# Distinct REV-ERBα Conformational State Predicted by GaMD Simulations Leads to the Structure-Based Discovery of Novel REV-ERBα Antagonist

**DOI:** 10.1101/2021.05.26.445894

**Authors:** Lamees Hegazy, Aurore-Cecile Valfort, Thomas P. Burris, Bahaa Elgendy

## Abstract

REV-ERBα is a nuclear hormone receptor that plays important role in the regulation of many physiological processes such as circadian clock regulation, inflammation, and metabolism. Despite its importance, few chemical tools are available to study this receptor. In addition, there is no available X-ray crystal structures of REV-ERB bound with synthetic ligands, hampering the development of targeted therapeutics. **SR8278** is the only identified synthetic antagonist of REV-ERB. We have performed Gaussian accelerated molecular dynamics (GaMD) simulations to sample the binding pathway of **SR8278** and associated conformational changes to REV-ERBα. The simulations revealed a novel and more energetically favorable conformational state than the starting conformation. The new conformation allows ligand binding to the orthosteric binding site in a specific orientation. This state is reached after a tryptophan (Trp436) rotameric switch coupled with H3-H6 distance change. We used the newly identified GaMD conformational state in structure-based virtual screening of one million compounds library which led to the identification of novel REV-ERBα antagonist. This study is the first that demonstrates a synthetic ligand binding pathway to REV-ERBα, which provided important insights into the REV-ERBα functional mechanism and lead to the discovery of novel REV-ERBα antagonists. This study further emphasizes the power of computational chemistry methods in advancing drug discovery research.

## Introduction

Nuclear receptors (NRs) constitute a superfamily of transcription factors that regulate gene transcription in response to various stimuli and control a myriad of biological and disease processes.^1,2^ Examples of well-known NRs are vitamin D receptor, retinoic acid receptor, and peroxisome proliferator-activated receptor. Nuclear hormone receptors (NRs) represent a major drug target, accounting for ~16% of all approved drugs.^3^ NRs have been targeted successfully in many therapeutic areas, including diabetes, skin disorders and breast and prostate cancers.^4–6^ A large number of NRs are referred to as ‘‘orphan’’ NRs because they don’t have identified natural ligands.^7^

NRs are characterized by multiple domain organization. Two of these domains are highly conserved and contribute to the activation of NRs: i. DNA-binding domain (DBD), located at the N terminus, known as activation function 1 (AF-1). DBD is ligand independent and interacts with target DNA sequences through two zinc fingers to recognize specific hormone response elements (HREs), ii. the Ligand binding domain (LBD), which is located at the C-terminal domain and interacts with small molecule ligands and cofactors involved in regulating the transcription, known as AF-2.^8^ The LBD is a globular domain composed almost exclusively of α-helices arranged as three layers “sandwich shape”. Ligands bind to the ligand binding pocket (LBP) within the interior of this globular domain and depending on the nature of the ligand, a conformational change of the LBD occurs leading to a cascade of downstream events. Agonists binding stabilize the helix12 orientation towards the ligand binding domain in a conformation favorable for coactivator binding, while binding of antagonists and inverse agonists cause displacement or structural rearrangement of helix12 that leads to interfering with the coactivator binding site.^9–13^

NRs assume a wide range of conformational states, including apo states, ligand specific states and recent studies implicate an important role for protein dynamics in the mechanism of action of nuclear receptor ligands.^15–17^ REV-ERBα and REV-ERBβ (encoded by NR1D1 and NR2D2, respectively) are a subfamily of the nuclear receptors (NRs). They are heme receptors and play important role in the regulation of many physiological processes such as inflammation and metabolism.^18,19^ REV-ERBs also play important role in the circadian clock regulation, memory and learning.^20–22^ Recent studies highlighted their role in neurological disorders such as cognitive diseases.^23^ Although REV-ERBs share similar molecular domain organization to most nuclear hormone receptors, they are distinctive because they lack the carboxy-terminal helix 12 activation function 2 (AF-2) region. Therefore, they are effective transcription repressors and interact constitutively with the NRs corepressors such as NCoR via helix 11 of the C-terminal ligand-binding domain.^24,25^ Upon binding to the DNA response element,^26^ REV-ERBs recruit corepressors to the target gene causing its repression through active histone deacetylation and condensation of the chromatin.^25,27^

The *in vivo* functions of REV-ERB have been widely explored over the past decade mostly through use of genetically engineered mice, but the lack of selective small molecule modulators hindered further exploration of this drug target in a more physiological context. Isserlin *et al*. analysed the citation patterns for NRs, and found that most of the research activity in this family is focused only on 10% of the family members and that ongoing research activities are influenced more by the availability of high quality chemical probes.^28^ The absence of essential structural information about the mechanism of ligand binding and distinct active and inactive conformations highly contributed to this problem. The heme bound REV-ERBβ is the only available X-ray structure available for ligand bound REV-ERB (Figure 1A).^24,29^ On the other hand, only one X-ray structure is reported for REV-ERBα LBD in which the receptor is crystallized bound with the corepressor NCoR ID1 (Figure 1B).^30^ The conformation of REV-ERB LBD in both structures has considerable differences specifically at the helix-3 and helix-11 regions. This information suggests that the LBP of REV-ERB is flexible and can accommodate different ligand scaffolds. There is a lack of chemical tools to study the biology of REV-ERB with only few ligands that have been used in the last decade. The most common ligands are **GSK4112**, **SR8278**, **SR9009**, and **SR9011**(Figure 2).^31^ These ligands suffer from high clearance rate and rapid metabolism and therefore display poor pharmacokinetics. Given this receptor pharmacological importance and clinical premise, understanding the molecular basis behind REV-ERB activation and ligand binding has profound implications for elucidating the detailed mechanism of this receptor and design of new therapeutic agents with subtype selectivity profile. **SR8278** is the only available antagonist to probe the function of REV-ERB in various disease models. ^32^ It suffers from low metabolic stability and short half-life (t1/2 = 0.17h). Identification of additional antagonists is necessary to investigate possible diverse signaling for REV-ERB in different disease models.

**Figure 1.**
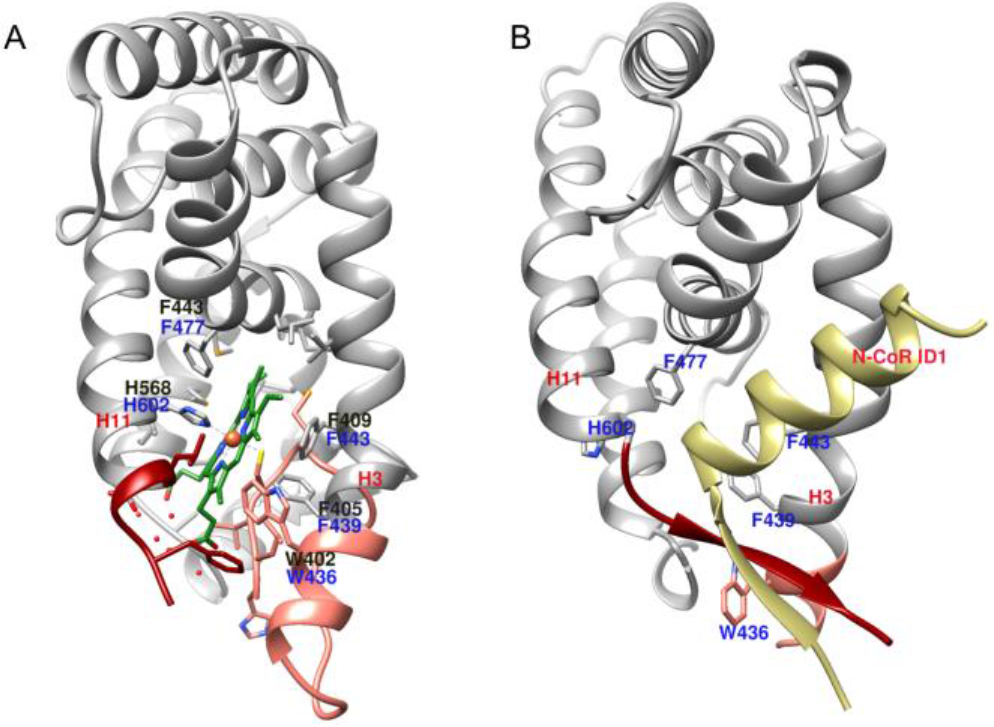
(A) X-ray structure of heme bound REVERBβ (PDB:3CQV). Residues labeled in black correspond to REV-ERBβ numbering while blue labels correspond to REV-ERBα numbering. (B) X-ray structure of REV-ERBα bound with the corepressor N-CoR ID1 (3N00).

**Figure 2.**
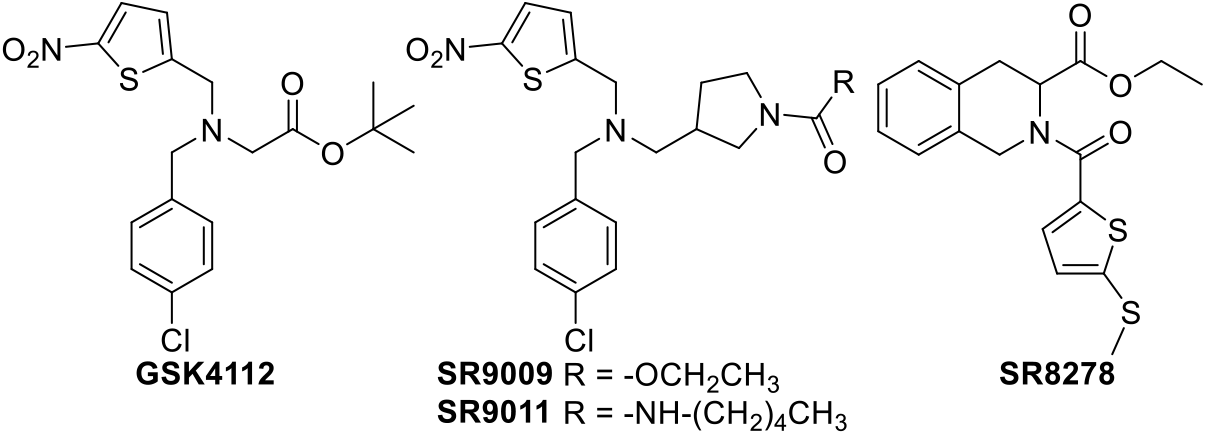
Chemical structure of available REV-ERB modulators.

There is no X-ray structure for REV-ERB bound with **SR2878**, therefore we used gaussian accelerated MD (GaMD) to predict ligand binding pathway of the REV-ERB antagonist (**SR8278**).^32^ GaMD have proven useful in revealing valuable information underlying ligand binding for different biological targets.^33–37^ The GaMD simulations revealed a novel conformational state that allows ligand binding to the orthosteric binding site in a specific orientation. We further used the newly identified GaMD conformational state in structure-based virtual screening of one million compounds library, which led to the identification of a novel REV-ERB antagonist. This study revealed the mechanism of **SR8278** binding to REV-ERBα, which in turn provided important insights into the REV-ERBα functional mechanism and enabled identification of novel REV-ERBα antagonists.

## RESULTS

To investigate the ligand binding pathway of the antagonist **SR8278** to REV-ERBα, we performed three independent molecular simulations (Table 1); one conventional molecular dynamics simulation (CMD) and two gaussian accelerated molecular dynamics simulations (GaMDs). In each simulation, 5 molecules of **SR8278** were placed in solvent at least 20 Å away from the protein (Figure S1). In the first CMD simulations, none of the **SR8278** molecules bind to the orthosteric pocket of REV-ERBα (Figure S2). However, one of the ligands was bound to the receptor surface near helix 6 for over 1 μs but it did not enter the orthosteric pocket and eventually dissociated into solvent (Figure S2F).

**Table 1.**

| ID | Simulation Type | Duration |
| --- | --- | --- |
| CMD | Classical MD | 3 μs |
| Sim1 | GAMD | 1 μs |
| Sim2 | GAMD | 700 ns |

In the other two GaMD simulations, ligand binding and dissociation to the orthosteric pocket was observed. In the first GaMD simulations (Sim1), ligand binding was observed after 100 ns, with repetitive ligand binding and dissociation events (Figure 3A). Although high flexibility was observed in the LBP region (helix 3, helix 6 and C-terminal domain of helix 11), no major conformational changes to the receptor were observed. The ligand entered the orthosteric pocket simultaneously through two entering points either through helix 3 and helix 5 or through helix 3 and helix 11 (Figure 3B). The ligand makes mainly hydrophobic interactions with amino acid residues F443, F439, M513 and L606 (Figure 3C).

**Figure 3.**
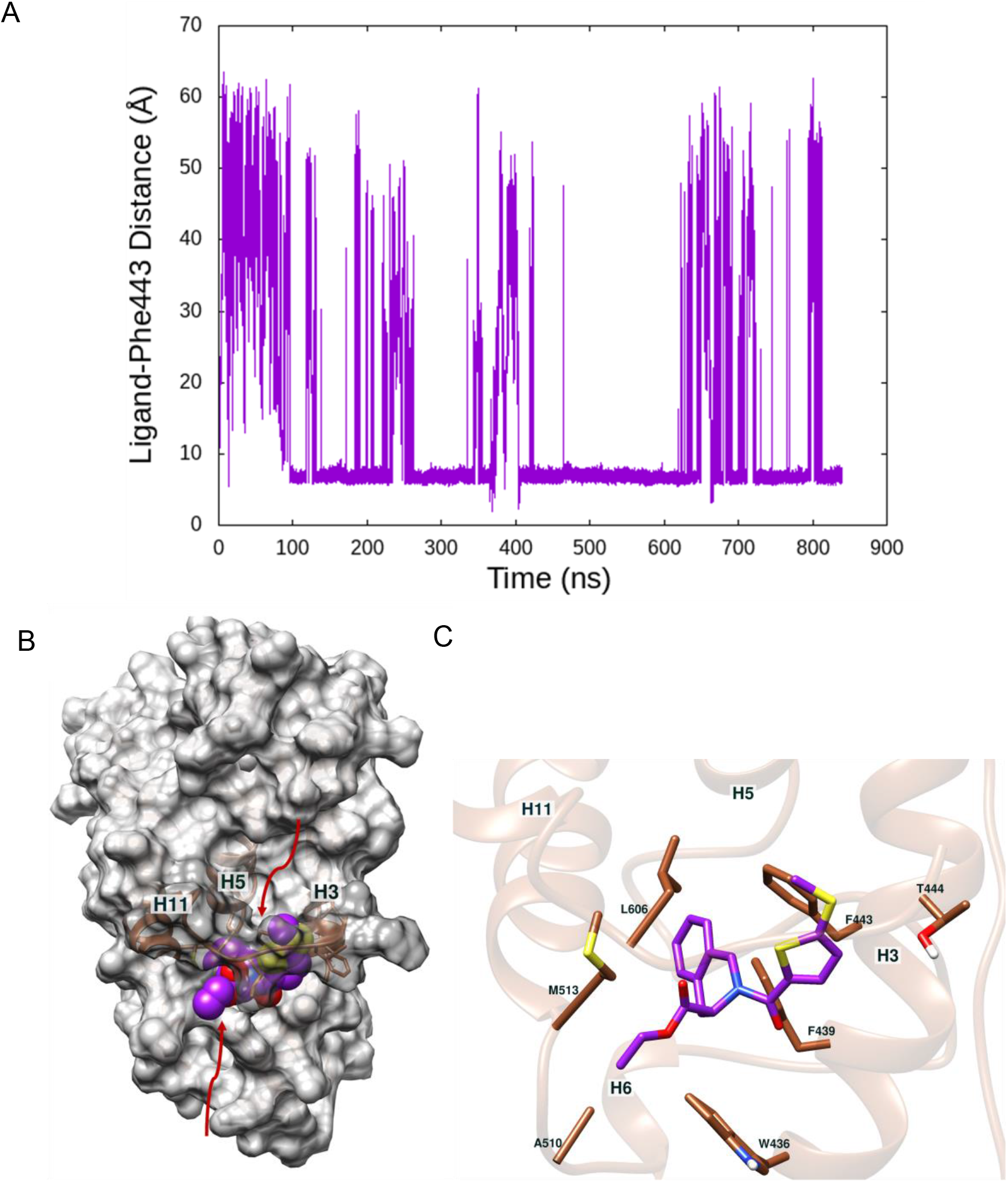
(A) Repetitive ligand binding and dissociation observed in GaMD (Sim1). (B) Ligand entrance pathways (red arrows). (C) Orientation and ligand binding pose in Sim1.

In the second GAMD simulations (Sim2), ligand binding was accompanied by major conformational changes where an increase in the distance between helix 3 and helix 6 occurred followed by upward shift of helix 3. At around 100 ns, the ligand initially made contacts with Trp436 at the exterior region between helix 6 and helix 3 for about 50 ns (Figure 4). At 150 ns, Trp436 changed its dihedral angle c-ca-cb-cg conformation from 50° to 180° (Fig. 5C). The free energy profile along this reaction coordinate shows that there are two minima for the Trp436 c-ca-cb-cg dihedral angle at 90° and 176° with the later value have more stable free energy (Figure 5D). The boost potential applied during the GAMD simulation follows Gaussian distribution and its distribution anharmonicity γ equals 0.013 suggesting efficient enhanced sampling (Figure S3). The region between H3 and H6 interchanged between two different states in absence and presence of the ligand. This conformational transition is monitored by the distance between H3 and H6 (Trp436-Ala126) (Figure 5A). The free energy profile along this reaction coordinate shows that in Sim1, only one energy minimum is observed at 10 Å, while in Sim2, ligand binding with different orientation induced distance shift between the two helices with a favourable free energy at around 13.5 Å (Figure 5B). The barrier between the two minima is 0.7 Kcal/mol thus suggesting ligand binding in orientation 2 (Sim2) induces distinctive minimum conformational state (Figure 5).

**Figure 4.**
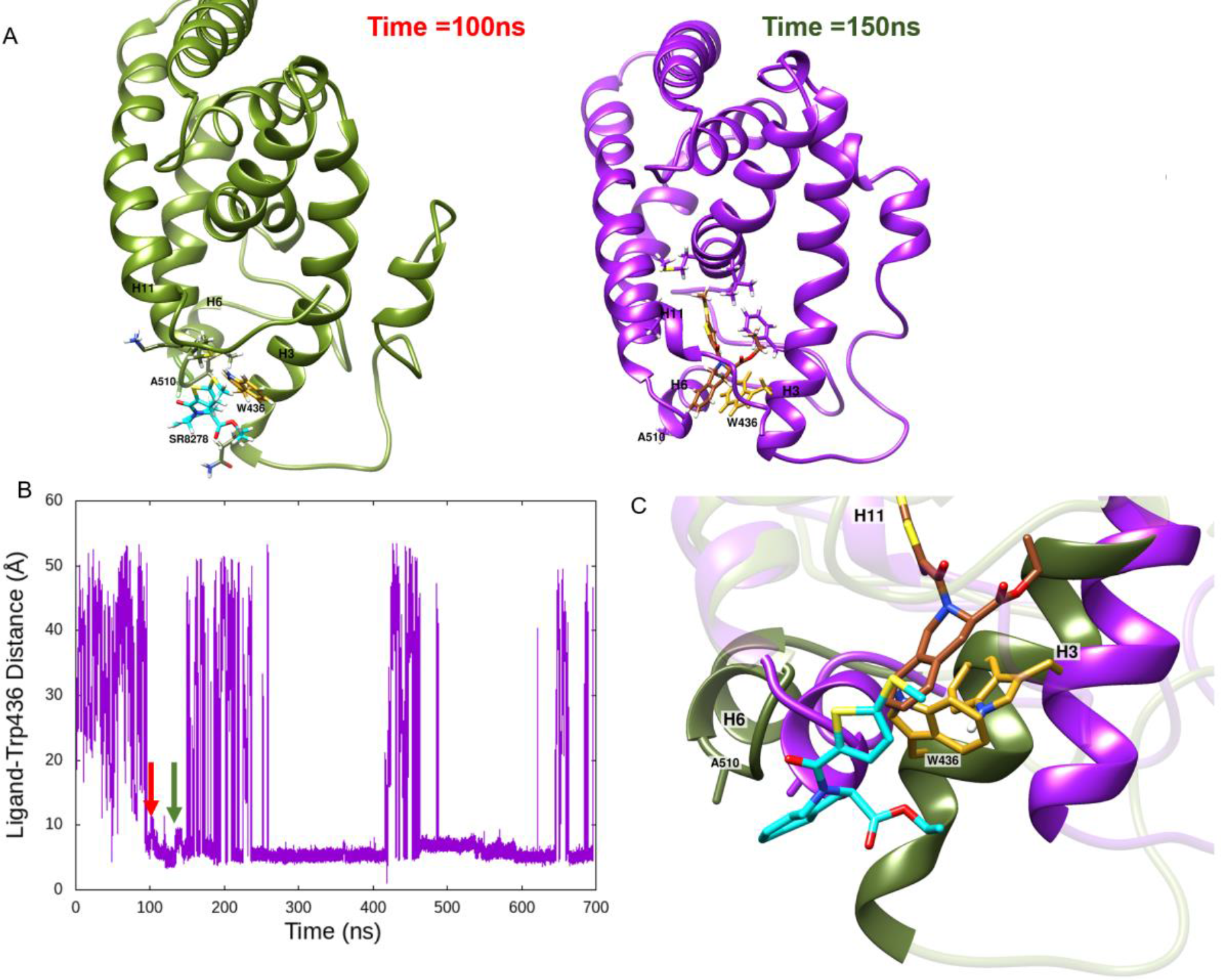
(A) Ligand entrance pathway and associated receptor conformational changes in Sim2. (B) Time course of the Ligand-Trp436 distance in Sim2 (C) Overlay and close view on ligand entrance into the ligand binding pocket at 100ns (red arrow) and 150ns (green arrow).

**Figure 5.**
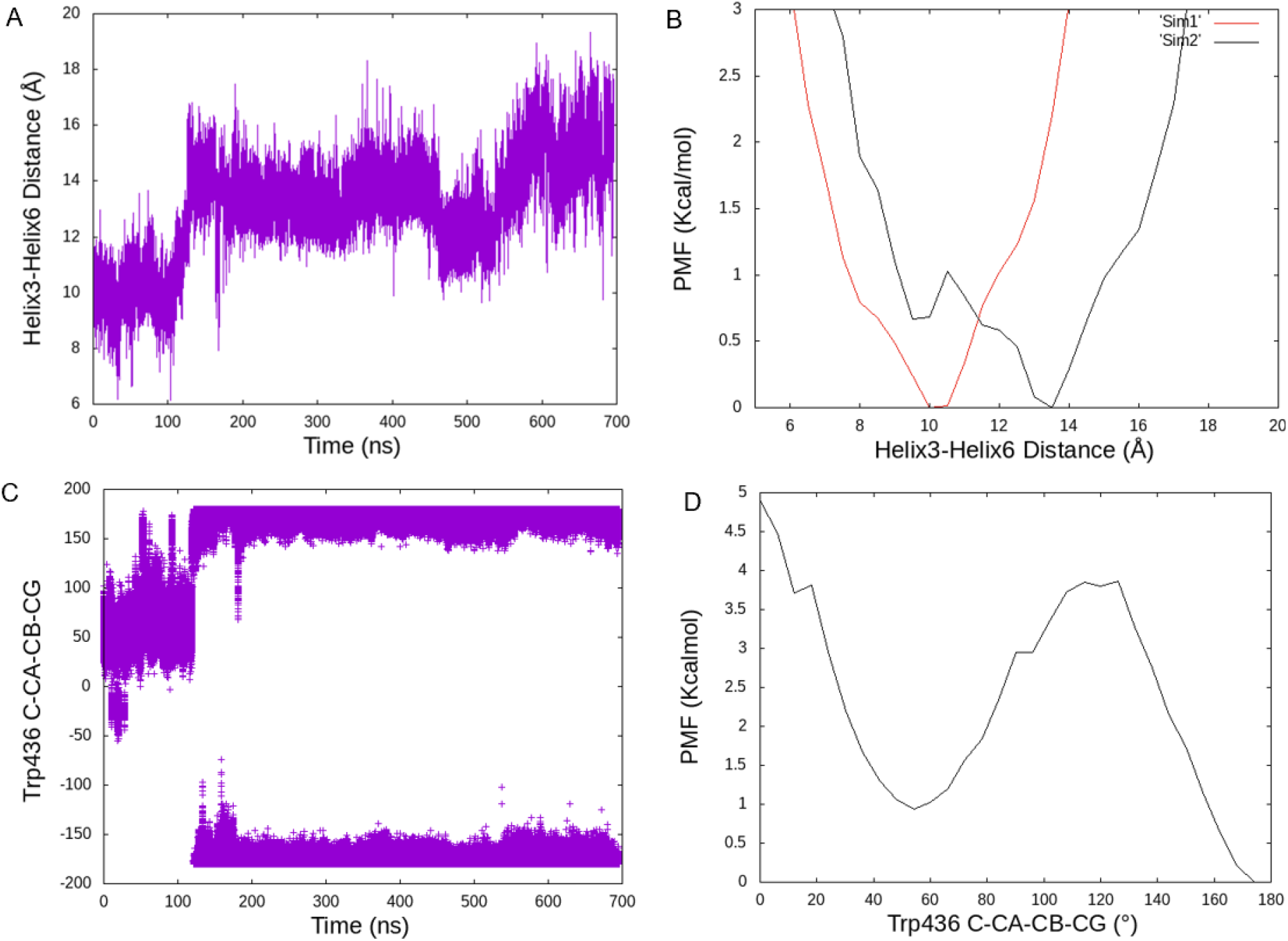
(A). Time course of helix3-helix6 distance in Sim2 (B) PMF profile calculated by the distance between helix 3 and helix 6 (C) Time course of Trp436 dihedral angle c-ca-cb-cg conformational change (D) PMF profile for Trp436 dihedral angle c-ca-cb-cg.

### Conventional Molecular Dynamics Simulations and Structure-based Virtual Screening

The prediction of the binding pose of **SR8278** with REV-ERBα provided us with an opportunity to seek previously undescribed antagonist chemotypes using structure-based virtual screening (SBVS). We first perofromed 500 ns of conventional molecular dynamics simulations on a snapshot from the ligand-bound GAMD simulations (Sim 2), to obtain an equilibrated state of the **SR8278**-bound REV-ERBα (Figure 6). **SR8278** is involved in hydrophobic interactions with amino acid residues F609, W436, F439, F484, L480, F477, L516 and L607. Seeking molecules that complemented the orthosteric site of the receptor, we docked more than one million ‘drug-like’ molecules, characterized by favourable physical properties from Enamine advance collection library. The best-scoring molecules were inspected for interactions with amino acid residues that were identified by the GaMD simulations to recognize **SR8278** and 19 compounds were selected for experimental testing in cotransfection cell based assay. We identified a novel chemotype as a lead compound, **BE7011** that has comparable potency to **SR8278** (**Table 2**). The identified REV-ERBα antagonist (i.e. **BE7011**) is a novel chemotype that agrees with Lipinski’s rule of 5 and is endowed with good physicochemical properties (MW<500, cLog P = 4.33, tPSA = 53.93, HBD = 1, and HBA = 4). To quantify the drug-likeness of our hit molecule, we used quantitative estimates of drug-likeness (QED) that is calculated based on the analysis of essential physiochemical properties: molecular weight, log P, number of hydrogen bond donors and acceptors, molecular polar surface area, number of rotatable bonds, number of aromatic rings, and number of structural alerts. This integrative approach weighs the contribution of main molecular properties according to their influence on the drug-likeness of the compound. QED value for our ligand is 0.69, which is above that of the top 30% of the ChEMBL database and it is optimal value for a starting hit molecule. **BE7011** showed evidence for structure activity in a follow-up screen of 3 commercially available analogs (i.e. **BE7201**, **BE7202**, and **BE7203**). Medicinal chemistry optimization of this lead series is under way to establish rigorous SAR and develop this new chemotype to REV-ERB antagonists that possess acceptable PK/PD properties.This study demonstrates the applicability of GaMD to the study of ligand binding pathway to nuclear receptors and identification of conformational states that is more suitable for exploitation by small-molecule inhibitors.

**Figure 6.**
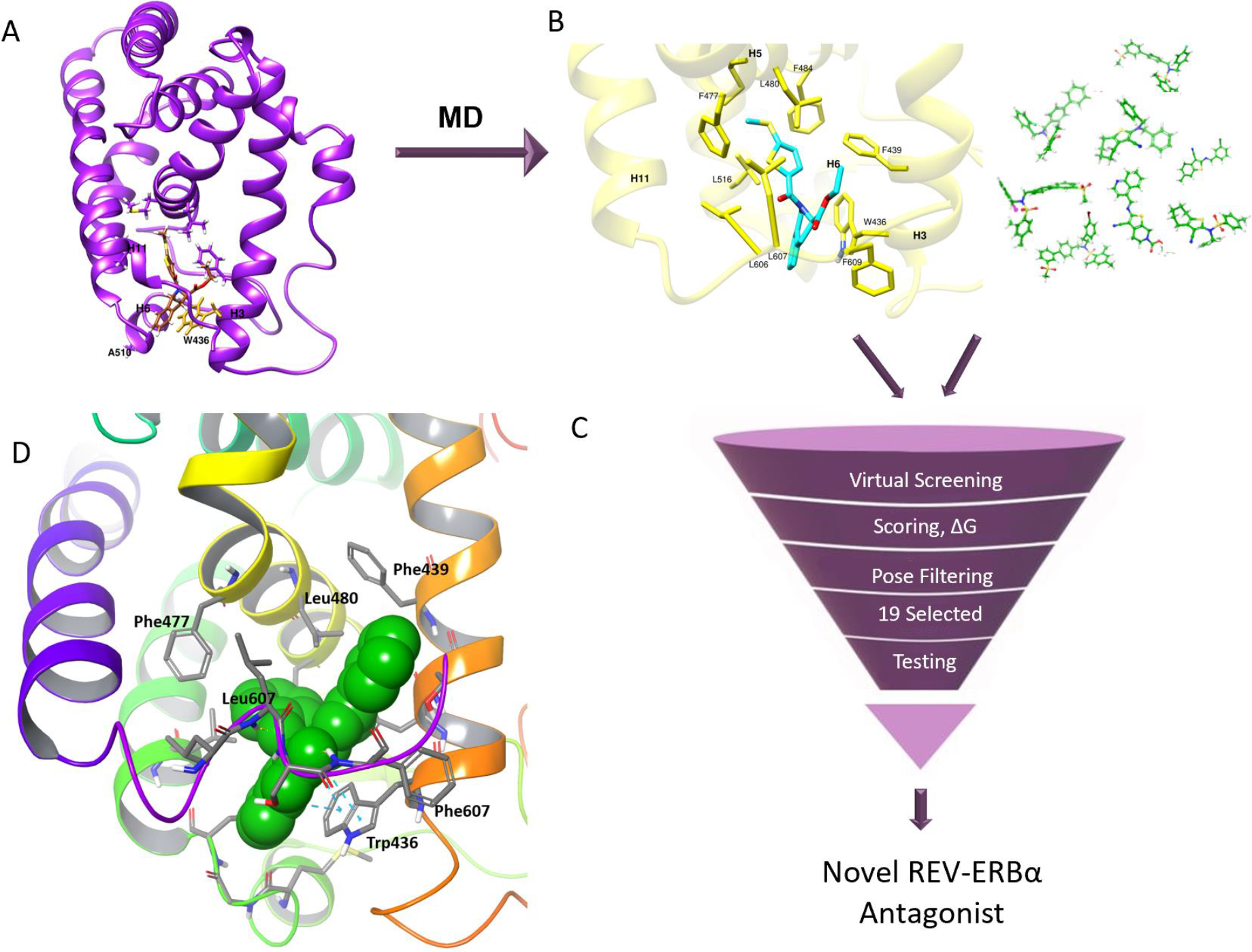
Virtual screening workflow for identification of novel REV-ERBα antagonist. (A) Snapshot from the GAMD simulations. (B) The binding pose of **SR8278** after CMD simulations and examples of compounds docked during the virtual screening. (C) Summary of the virtual screening and filtering workflow. (D) Docking pose of the novel antagonist **BE7011**

**Table 2.** EC_50_ values for REV-ERB antagonists identified by virtual screening.

| Compound | EC<sub>50</sub> (μM) |
| --- | --- |
| **SR828** | 2.267 |
| **BE7011** | 2.724 |
| **BE7201** | 2.599 |
| **BE7202** | 2.768 |
| **BE7203** | 2.803 |

**Figure 7.**
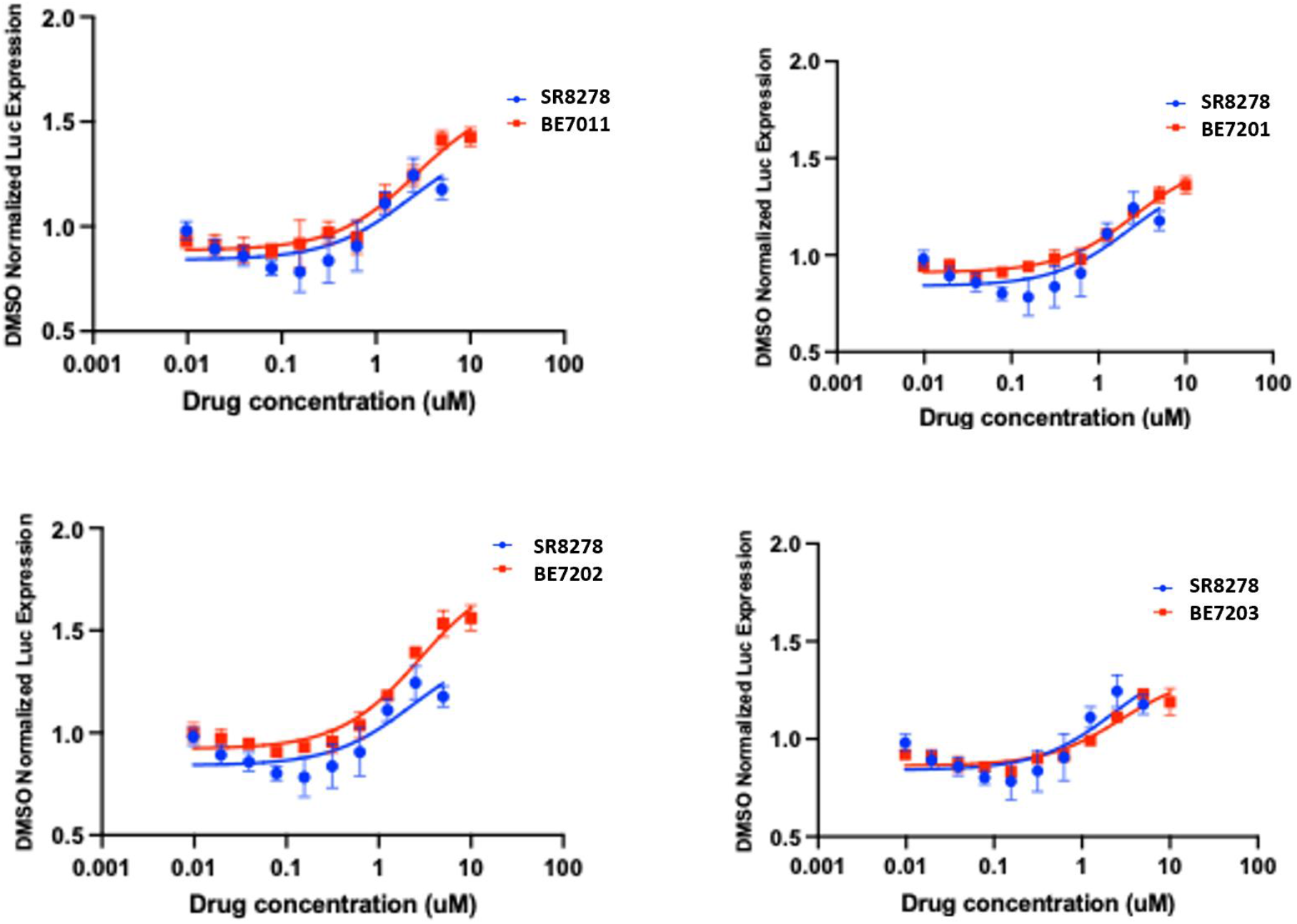
Functional characterization of the virtual screening hit **BE7011** and its analogs for antagonist activity at REV-ERBα.

## Discussion

In this study we investigated the ligand binding mechanism of **SR8278** to REV-ERBα using GaMD simulations. GaMD simulations captured complete binding of the antagonist **SR8278** to the ligand-binding pocket of REV-ERBα. Two different orientations were predicted with low-energy bound states from the reweighted free energy profiles (Sim1 & Sim2). Ligand binding orientation in Sim2 induced distinct conformational state initiated by Trp436 rotemeirc switch, upward shift of helix3, and distance increase between helix3 and helix6. This state has a favorable binding energy than the conformational state identified in Sim1. To confirm the applicability of newly identified conformational state in drug discovery, we carried out *in silico* virtual screening of one million compounds from the Enamine Advanced library in the conformational state identified by Sim2. A novel REV-ERBα antagonist, **BE7011** was identified and confirmed in cell-based cotransfection assay. **BE7011** showed evidence for structure activity in a follow-up screen of 3 commercially available analogs. Medicinal chemistry optimization of this lead series is under way to establish rigorous SAR and develop this new chemotype to REV-ERB antagonists that possess acceptable PK/PD properties.This is the first study that reveals **SR8278** binding mechanism and identify novel REV-ERBα antagonist through structure-based virtual screening using a conformational ensembel generated by molecular simulations. This study emphasizes the importance of integrating computer-based methods for accelerating drug discovery research.

## METHODS

### GaMD Simulations Set up & Analysis

GaMD simulations were performed on the REV-ERBα Xray structure bound with corepressor NCoR (PDB: 3N00).^30^ The corepressor was removed during set up of the simulations. All mutations and gaps were fixed and missing residues 155-183 were modelled using I-TASSER suite (Loop T155-E179).^38,39^ Ligand parameters were assigned according to the general AMBER force field (GAFF) and the corresponding AM1BCC charges using Antechamber.^40^ The FF14SB forcefield parameters were used for all receptor residues.^41^ Tleap module was used to neutralize and solvate the complexes using an octahedral water box of TIP3P water molecules. Simulations were performed on GPUs in Amber16 using the CUDA version of Particle Mesh Ewald Molecular Dynamics (PMEMD).^42^ Each system was first energy minimized using the steepest descent and conjugate gradient methods. Then gradually heated with the Langevin thermostat to 300K over 30 PS at constant volume using 1fs time step. Initial velocities were sampled from the Boltzman distribution while keeping week restraints on the solute and the ligand. Each system was then equilibrated in the isothermal-isobaric ensemble (NPT), at 300 K, using constant pressure periodic boundary with an average pressure of 1 atm. Isotropic position scaling was used to maintain the pressure with a relaxation time of 2 ps. Non-bonded interactions were cut off at 8.0 A° and long-range electrostatic interactions were computed using the particle mesh Ewald (PME). The SHAKE algorithm was used to keep bonds involving H atoms at their equilibrium length. 2 fs time step was used for the integration of Newton’s equations. GaMD simulation were performed using the GaMD module implemented in the graphics processing unit (GPU) version of AMBER18.^33,35^The simulations started with 10-ns short cMD simulation used to collect potential statistics for calculating the GaMD acceleration parameters followed by 50-ns equilibration after adding the boost potential, and finally multiple independent GaMD production runs with randomized initial atomic velocities. All GaMD simulations were run at the “dual-boost”: one boost potential is applied to the dihedral energetic term and another to the total potential energetic term. The threshold energy was set to the lower bound E =Vmax. The average and standard deviation (SD) of the system potential energies were calculated every 200,000 steps (400 ps). The upper limit of the boost potential SD, σ0 was set to 6.0 kcal/mol for both the dihedral and the total potential energetic terms. CPPTRAJ and Chimera were used to analyze the GaMD simulation trajectories.^43,44^ The free energy profile was calculated using cumulant expansion to the first order with the PyReweighting toolkit.^45^

### Structure-based Virtual Screening

A snapshot of ligand bound REV-ERBα from the GaMD simulations was first subjected to conventional molecular dynamics (cMD) simulations to obtain equilibrated conformational ensemble. The conventional cMD simulations were performed using a similar protocol to the cMD equilibration run (before the GaMD equilibration & production runs) described in the section above. The virtual screening calculations was accomplished using the Schrodinger suite for small molecule docking.^46^ The structure was first prepared for docking using the Schrodinger Protein Prep Wizard by adding missing hydrogens, assigning side chain protonation states, and refining the structure (energy minimization). The receptor grid generation tool was used to generate receptor grid that represent the shape and properties of the receptor. We used the Enamine advanced collection library in the virtual screening. The ligands were initially prepared using the LigPrep tool implemented in Schrodinger to correct tautomeric and ionization states, ring conformations, and stereoisomers to produce broad chemical and structural diversity from a single input structure. The docking calculations were performed using Glide standard precision (SP) scoring function.^46^ Computations were performed at the Center for High Performance Computing, Mallinckrodt Institute of Radiology, Washington University in Saint Louis.

### Cell Culture and Cotransfections

HEK293T cells (ATCC) were maintained in Dulbecco’s modified Eagle’s medium (DMEM) supplemented with 10% fetal bovine serum at 37°C under 5%CO_2_. Twenty-four hours prior to drug treatment, cells were reverse transfected and plated in 96-well plates at a density of 3.5×104 cells/well. Transfections were performed using Lipofectamine 2000 (Invitrogen). Cells were treated with vehicle (DMSO) or compound. Twenty-four hours post-treatment, the luciferase activity was measured using the One-Glo luciferase assay system (Promega). The values indicated represent the means and standard errors from four independently transfected wells. The REV-ERBR and reporter constructs have been previously described.^32^

## ACKNOWLEDGMENTS

The authors would like to thank Professor Yinglong Miao at University of Kansas for valuable discussions. We are grateful to the funding provided by the Department of Defense office of the Congressionally Directed Medical Research Programs (CDMRP in support of this project (W81XWH-19-1-0633).

## References

(1) Mangelsdorf, D. J.; Thummel, C.; Beato, M.; Herrlich, P.; Schütz, G.; Umesono, K.; Blumberg, B.; Kastner, P.; Mark, M.; Chambon, P.; Evans, R. M. The Nuclear Receptor Superfamily: The Second Decade. Cell 1995, 83 (6), 835–839. https://doi.org/10.1016/0092-8674(95)90199-x.

(2) Bookout, A. L.; Jeong, Y.; Downes, M.; Yu, R. T.; Evans, R. M.; Mangelsdorf, D. J. Anatomical Profiling of Nuclear Receptor Expression Reveals a Hierarchical Transcriptional Network. Cell 2006, 126 (4), 789–799. https://doi.org/10.1016/j.cell.2006.06.049.

(3) Santos, R.; Ursu, O.; Gaulton, A.; Bento, A. P.; Donadi, R. S.; Bologa, C. G.; Karlsson, A.; Al-Lazikani, B.; Hersey, A.; Oprea, T. I.; Overington, J. P. A Comprehensive Map of Molecular Drug Targets. Nat. Rev. Drug Discov. 2017, 16 (1), 19–34. https://doi.org/10.1038/nrd.2016.230.

(4) Schulman, I. G. Nuclear Receptors as Drug Targets for Metabolic Disease. 2011, 62 (13), 1307–1315. https://doi.org/10.1016/j.addr.2010.07.002.Nuclear.

(5) Burris, T. P.; Solt, L. A.; Wang, Y.; Crumbley, C.; Banerjee, S.; Griffett, K.; Lundasen, T.; Hughes, T.; Kojetin, D. J. Nuclear Receptors and Their Selective Pharmacologic Modulators. Pharmacol. Rev. 2013, 65 (2), 710–778. https://doi.org/10.1124/pr.112.006833.

(6) Yin, K.; Smith, A. G. Nuclear Receptor Function in Skin Health and Disease: Therapeutic Opportunities in the Orphan and Adopted Receptor Classes. Cellular and Molecular Life Sciences. Springer International Publishing 2016, pp 3789–3800. https://doi.org/10.1007/s00018-016-2329-4.

(7) Mullican, S. E.; Dispirito, J. R.; Lazar, M. A. The Orphan Nuclear Receptors at Their 25-Year Reunion. J. Mol. Endocrinol. 2013, 51 (3), T115–T140. https://doi.org/10.1530/JME-13-0212.

(8) Novac, N.; Heinzel, T. Nuclear Receptors: Overview and Classification. Curr. Drug Targets. Inflamm. Allergy 2004, 3 (4), 335–346. https://doi.org/10.2174/1568010042634541.

(9) Meijer, F. A.; Leijten-van de Gevel, I. A.; de Vries, R. M. J. M.; Brunsveld, L. Allosteric Small Molecule Modulators of Nuclear Receptors. Mol. Cell. Endocrinol. 2019, 485, 20–34. https://doi.org/10.1016/j.mce.2019.01.022.

(10) Helsen, C.; Claessens, F. Looking at Nuclear Receptors from a New Angle. Mol. Cell. Endocrinol. 2014, 382 (1), 97–106. https://doi.org/10.1016/j.mce.2013.09.009.

(11) Jin, L.; Li, Y. Structural and Functional Insights into Nuclear Receptor Signaling. Adv. Drug Deliv. Rev. 2010, 62 (13), 1218–1226. https://doi.org/10.1016/j.addr.2010.08.007.

(12) Rastinejad, F.; Huang, P.; Chandra, V.; Khorasanizadeh, S. Understanding Nuclear Receptor Form and Function Using Structural Biology. J. Mol. Endocrinol. 2013, 51 (3), T1–T21. https://doi.org/10.1530/JME-13-0173.

(13) Moras, D.; Gronemeyer, H. The Nuclear Receptor Ligand-Binding Domain: Structure and Function. Curr. Opin. Cell Biol. 1998, 10 (3), 384–391. https://doi.org/10.1016/S0955-0674(98)80015-X.

(14) Flaveny, C. A.; Solt, L. A.; Kojetin, D. J.; Burris, T. P. Chapter 4 - Biased Signaling and Conformational Dynamics in Nuclear Hormone Receptors; Arey Pharmacology and Therapeutics, B. J. B. T.-B. S. in P., Ed.; Academic Press: San Diego, 2014; pp 103–135. https://doi.org/10.1016/B978-0-12-411460-9.00004-5.

(15) Hughes, T. S.; Chalmers, M. J.; Novick, S.; Kuruvilla, D. S.; Chang, M. R.; Kamenecka, T. M.; Rance, M.; Johnson, B. A.; Burris, T. P.; Griffin, P. R.; Kojetin, D. J. Ligand and Receptor Dynamics Contribute to the Mechanism of Graded PPARγ Agonism. Structure 2012, 20 (1), 139–150. https://doi.org/10.1016/j.str.2011.10.018.

(16) Hughes, T. S.; Shang, J.; Brust, R.; de Vera, I. M. S.; Fuhrmann, J.; Ruiz, C.; Cameron, M. D.; Kamenecka, T. M.; Kojetin, D. J. Probing the Complex Binding Modes of the PPARγ Partial Agonist 2-Chloro-N-(3-Chloro-4-((5-Chlorobenzo[d]Thiazol-2-Yl)Thio)Phenyl)-4-(Trifluoromethyl)Benzenesulfonamide (T2384) to Orthosteric and Allosteric Sites with NMR Spectroscopy. J. Med. Chem. 2016, 59 (22), 10335–10341. https://doi.org/10.1021/acs.jmedchem.6b01340.

(17) Kojetin, D. J.; Burris, T. P. Small Molecule Modulation of Nuclear Receptor Conformational Dynamics: Implications for Function and Drug Discovery. Mol. Pharmacol. 2013, 83 (1), 1–8. https://doi.org/10.1124/mol.112.079285.

(18) Burris, T. P. Nuclear Hormone Receptors for Heme: REV-ERBalpha and REV-ERBbeta Are Ligand-Regulated Components of the Mammalian Clock. Mol. Endocrinol. 2008, 22 (7), 1509–1520. https://doi.org/10.1210/me.2007-0519.

(19) Raghuram, S.; Stayrook, K. R.; Huang, P.; Rogers, P. M.; Nosie, A. K.; McClure, D. B.; Burris, L. L.; Khorasanizadeh, S.; Burris, T. P.; Rastinejad, F. Identification of Heme as the Ligand for the Orphan Nuclear Receptors REV-ERBalpha and REV-ERBbeta. Nat. Struct. Mol. Biol. 2007, 14 (12), 1207–1213. https://doi.org/10.1038/nsmb1344.

(20) Solt, L. A.; Wang, Y.; Banerjee, S.; Hughes, T.; Kojetin, D. J.; Lundasen, T.; Shin, Y.; Liu, J.; Cameron, M. D.; Noel, R.; Yoo, S.-H.; Takahashi, J. S.; Butler, A. A.; Kamenecka, T. M.; Burris, T. P. Regulation of Circadian Behaviour and Metabolism by Synthetic REV-ERB Agonists. Nature 2012, 485 (7396), 62–68. https://doi.org/10.1038/nature11030.

(21) Griffin, P.; Dimitry, J. M.; Sheehan, P. W.; Lananna, B. V; Guo, C.; Robinette, M. L.; Hayes, M. E.; Cedeño, M. R.; Nadarajah, C. J.; Ezerskiy, L. A.; Colonna, M.; Zhang, J.; Bauer, A. Q.; Burris, T. P.; Musiek, E. S. Circadian Clock Protein Rev-Erbα Regulates Neuroinflammation. Proc. Natl. Acad. Sci. 2019, 116 (11), 5102–5107. https://doi.org/10.1073/pnas.1812405116.

(22) Crumbley, C.; Burris, T. P. Direct Regulation of CLOCK Expression by REV-ERB. PLoS One 2011, 6 (3), e17290.

(23) Alzheimer’s Association. 2018 Alzheimer’s Disease Facts and Figures. Alzheimers Dement 2018;14(3):367–429; 2018. https://doi.org/10.1016/j.jalz.2009.03.001.

(24) Woo, E.-J.; Jeong, D. G.; Lim, M.-Y.; Jun Kim, S.; Kim, K.-J.; Yoon, S.-M.; Park, B.-C.; Eon Ryu, S. Structural Insight into the Constitutive Repression Function of the Nuclear Receptor Rev-Erbβ. J. Mol. Biol. 2007, 373 (3), 735–744. https://doi.org/10.1016/j.jmb.2007.08.037.

(25) Yin, L.; Lazar, M. A. The Orphan Nuclear Receptor Rev-Erbα Recruits the N-CoR/Histone Deacetylase 3 Corepressor to Regulate the Circadian Bmal1 Gene. Mol. Endocrinol. 2005, 19 (6), 1452–1459. https://doi.org/10.1210/me.2005-0057.

(26) Sierk, M. L.; Zhao, Q.; Rastinejad, F. DNA Deformability as a Recognition Feature in the RevErb Response Element. Biochemistry 2001, 40 (43), 12833–12843. https://doi.org/10.1021/bi011086r.

(27) Harding, H. P.; Lazar, M. A. The Monomer-Binding Orphan Receptor Rev-Erb Represses Transcription as a Dimer on a Novel Direct Repeat. Mol. Cell. Biol. 1995, 15 (9), 4791–4802. https://doi.org/10.1128/mcb.15.9.4791.

(28) Isserlin, R.; Bader, G.; Edwards, A.; Frye, S.; Willson, T.; Yu, F. The Human Genome and Drug Discovery after a Decade. Roads (Still) Not Taken. 2011.

(29) Pardee, K. I.; Xu, X.; Reinking, J.; Schuetz, A.; Dong, A.; Liu, S.; Zhang, R.; Tiefenbach, J.; Lajoie, G.; Plotnikov, A. N.; Botchkarev, A.; Krause, H. M.; Edwards, A. The Structural Basis of Gas-Responsive Transcription by the Human Nuclear Hormone Receptor REV-ERBbeta. PLoS Biol. 2009, 7 (2), e43–e43. https://doi.org/10.1371/journal.pbio.1000043.

(30) Phelan, C. A.; Gampe Jr, R. T.; Lambert, M. H.; Parks, D. J.; Montana, V.; Bynum, J.; Broderick, T. M.; Hu, X.; Williams, S. P.; Nolte, R. T.; Lazar, M. A. Structure of Rev-Erbα Bound to N-CoR Reveals a Unique Mechanism of Nuclear Receptor–Co-Repressor Interaction. Nat. Struct. & Mol. Biol. 2010, 17, 808.

(31) Kojetin, D. J.; Burris, T. P. REV-ERB and ROR Nuclear Receptors as Drug Targets. Nat. Rev. Drug Discov. 2014, 13 (3), 197–216. https://doi.org/10.1038/nrd4100.

(32) Kojetin, D.; Wang, Y.; Kamenecka, T. M.; Burris, T. P. Identification of SR8278, a Synthetic Antagonist of the Nuclear Heme Receptor REV-ERB. ACS Chem. Biol. 2011, 6 (2), 131–134. https://doi.org/10.1021/cb1002575.

(33) Miao, Y.; McCammon, J. A. Chapter Six - Gaussian Accelerated Molecular Dynamics: Theory, Implementation, and Applications; Dixon, D. A. B. T.-A. R. in C. C., Ed.; Elsevier, 2017; Vol. 13, pp 231–278. https://doi.org/10.1016/bs.arcc.2017.06.005.

(34) Miao, Y.; McCammon, J. A. Graded Activation and Free Energy Landscapes of a Muscarinic G-Protein–Coupled Receptor. Proc. Natl. Acad. Sci. 2016, 113 (43), 12162 LP –12167. https://doi.org/10.1073/pnas.1614538113.

(35) Bhattarai, A.; Miao, Y. Gaussian Accelerated Molecular Dynamics for Elucidation of Drug Pathways. Expert Opin. Drug Discov. 2018, 13 (11), 1055–1065. https://doi.org/10.1080/17460441.2018.1538207.

(36) An, X.; Bai, Q.; Bing, Z.; Zhou, S.; Shi, D.; Liu, H.; Yao, X. How Does Agonist and Antagonist Binding Lead to Different Conformational Ensemble Equilibria of the κ-Opioid Receptor: Insight from Long-Time Gaussian Accelerated Molecular Dynamics Simulation. ACS Chem. Neurosci. 2019, 10 (3), 1575–1584. https://doi.org/10.1021/acschemneuro.8b00535.

(37) Kappel, K.; Miao, Y.; McCammon, J. A. Accelerated Molecular Dynamics Simulations of Ligand Binding to a Muscarinic G-Protein-Coupled Receptor. Q. Rev. Biophys. 2015, 48 (4), 479–487. https://doi.org/10.1017/S0033583515000153.

(38) Zhang, Y. I-TASSER Server for Protein 3D Structure Prediction. BMC Bioinformatics 2008, 9 (1), 40. https://doi.org/10.1186/1471-2105-9-40.

(39) Yang, J.; Yan, R.; Roy, A.; Xu, D.; Poisson, J.; Zhang, Y. The I-TASSER Suite: Protein Structure and Function Prediction. Nat. Methods 2015, 12 (1), 7–8. https://doi.org/10.1038/nmeth.3213.

(40) Kumar, S.; Rosenberg, J. M.; Bouzida, D.; Swendsen, R. H.; Kollman, P. A. Multidimensional Free-Energy Calculations Using the Weighted Histogram Analysis Method. J. Comput. Chem. 1995, 16 (11), 1339–1350. https://doi.org/10.1002/jcc.540161104.

(41) Ponder, J. W.; Case, D. A. Force Fields for Protein Simulations. 2003, 66, 27–85.

(42) Salomon-Ferrer, R.; Case, D. A.; Walker, R. C. An Overview of the Amber Biomolecular Simulation Package. Wiley Interdiscip. Rev. Comput. Mol. Sci. 2013, 3 (2), 198–210. https://doi.org/10.1002/wcms.1121.

(43) Roe, D. R.; Cheatham, T. E. PTRAJ and CPPTRAJ: Software for Processing and Analysis of Molecular Dynamics Trajectory Data. J. Chem. Theory Comput. 2013, 9 (7), 3084–3095. https://doi.org/10.1021/ct400341p.

(44) Pettersen, E. F.; Goddard, T. D.; Huang, C. C.; Couch, G. S.; Greenblatt, D. M.; Meng, E. C.; Ferrin, T. E. UCSF Chimera—A Visualization System for Exploratory Research and Analysis. J. Comput. Chem. 2004, 25 (13), 1605–1612. https://doi.org/10.1002/jcc.20084.

(45) Miao, Y.; Sinko, W.; Pierce, L.; Bucher, D.; Walker, R. C.; McCammon, J. A. Improved Reweighting of Accelerated Molecular Dynamics Simulations for Free Energy Calculation. J. Chem. Theory Comput. 2014, 10 (7), 2677–2689. https://doi.org/10.1021/ct500090q.

(46) Friesner, R. A.; Banks, J. L.; Murphy, R. B.; Halgren, T. A.; Klicic, J. J.; Mainz, D. T.; Repasky, M. P.; Knoll, E. H.; Shelley, M.; Perry, J. K.; Shaw, D. E.; Francis, P.; Shenkin, P. S. Glide: A New Approach for Rapid, Accurate Docking and Scoring. 1. Method and Assessment of Docking Accuracy. J. Med. Chem. 2004, 47 (7), 1739–1749. https://doi.org/10.1021/jm0306430.

