## supplemental supporting information for "Distinct REV-ERBα Conformational State Predicted by GaMD Simulations Leads to the Structure-Based Discovery of Novel REV-ERBα Antagonist"

### Supplementary Figures

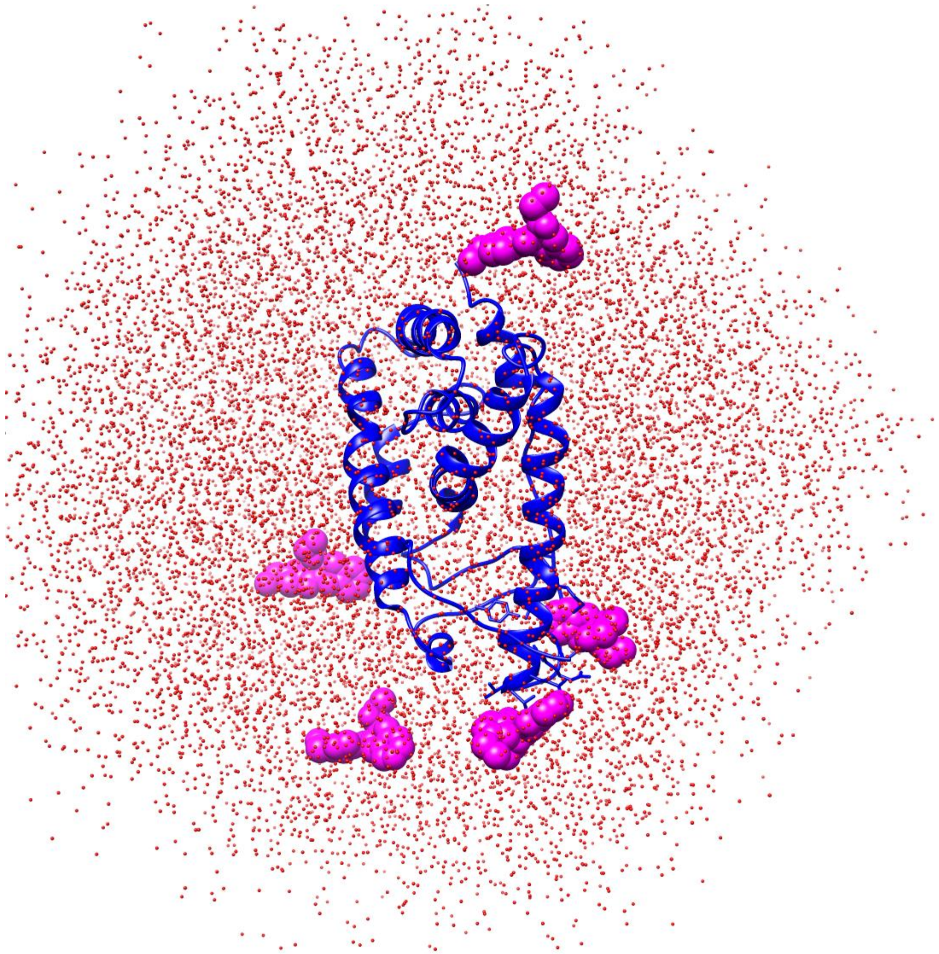

**Figure S1**. Initial model used in the GaMD simulations. REV-ERBα shown as blue ribbons and **SR8278** is shown as pink sphere representation.

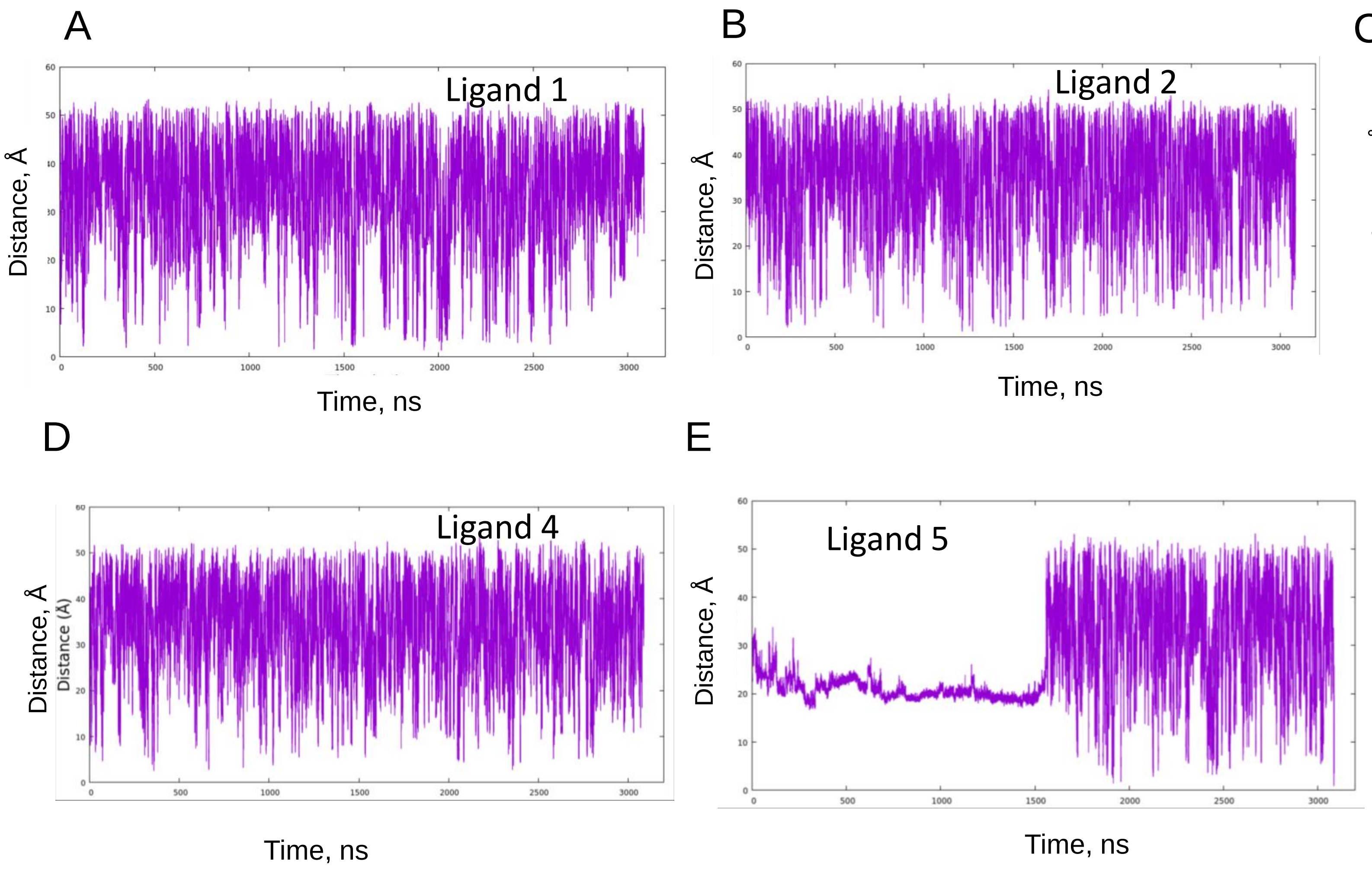


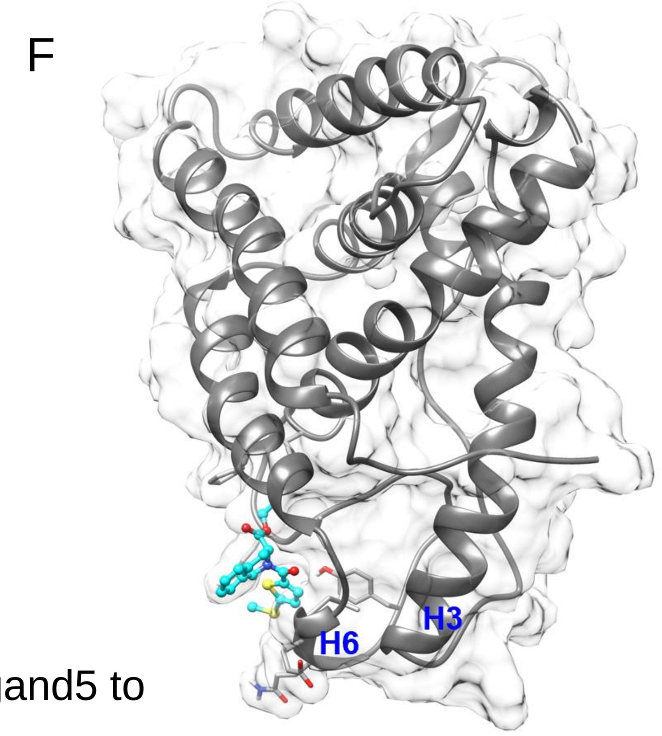


Figure S2-1. (A-E) Ligand-Trp436 Distance vs. simulation time plots. F. Binding of Ligand5 to the surface near helix 6 before dissociation.

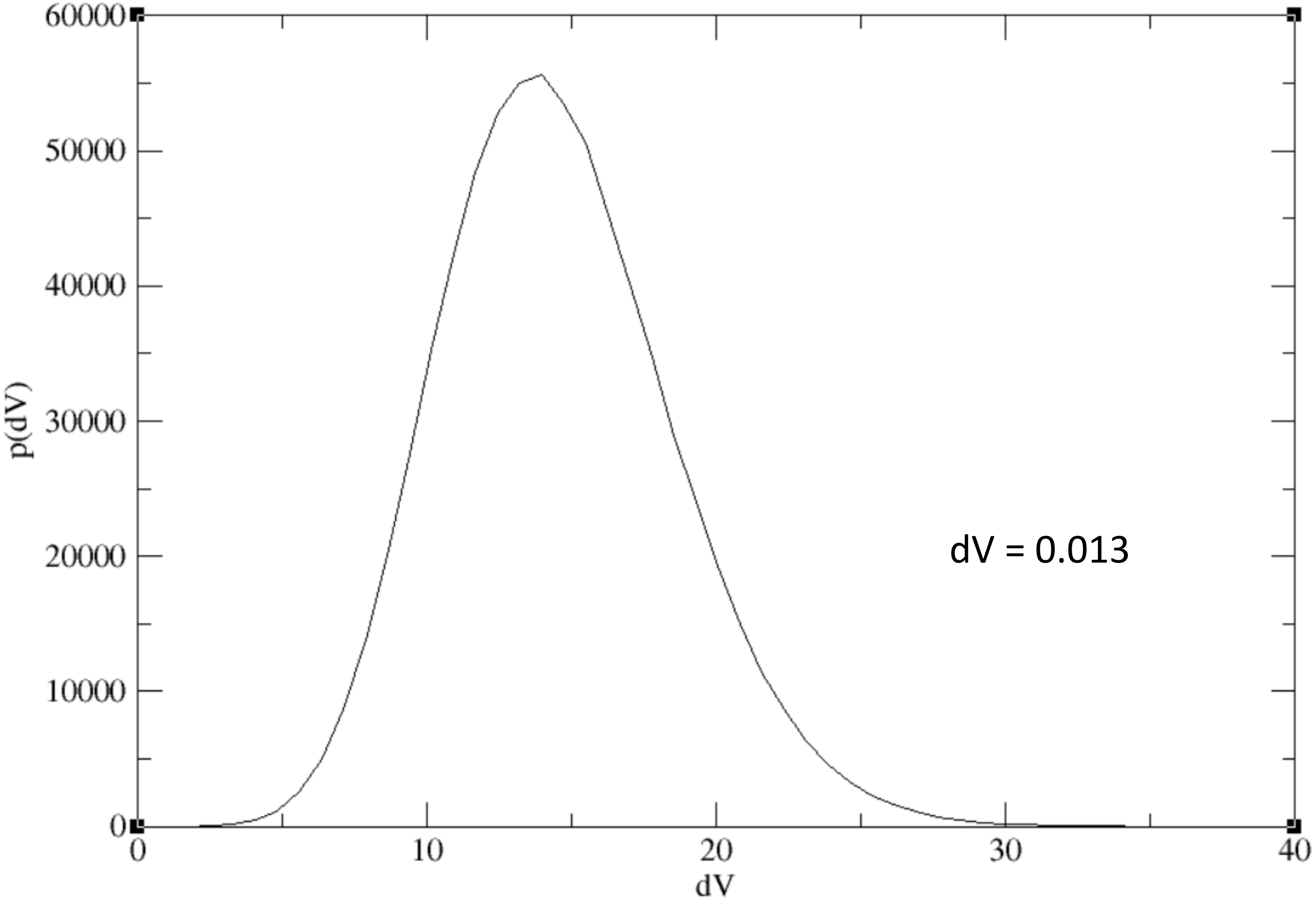


Figure S3. Distribution of the boost potential ΔV
